# Cost and benefits of being social: examining the influence of sociality on faecal parasites in free-ranging rhesus macaques

**DOI:** 10.1101/2022.12.20.521230

**Authors:** Melissa A. Pavez-Fox, Carla M. Escabi-Ruiz, Jordan D. A. Hart, Josue E. Negron-Del Valle, Daniel Phillips, Michael J. Montague, Michael L. Platt, Angelina Ruiz-Lambides, Melween I. Martinez, Cayo Biobank Research Unit, James P. Higham, Noah Snyder-Mackler, Lauren J.N. Brent

## Abstract

Parasites and infectious diseases constitute an important challenge to the health of group-living animals. Social contact and shared space can both increase disease transmission risk, while individual differences in social resources can help prevent infections. For example, high social status individuals and those with more or stronger social relationships may have better immunity and, thus, lower parasitic burden. To test for health trade-offs in the costs and benefits of sociality, we quantified how parasitic load varied with an individual’s social status, as well as with their weak and strong affiliative relationships in a free-ranging population of rhesus macaques (*Macaca mulatta*). Social resources may also protect against infection under environmentally challenging situations, such as natural disasters. We additionally examined the impact of a major hurricane on the sociality-parasite relationship in this system. We found that both weak and strong proximity partners, but not grooming partners, were associated with lower protozoa infection risk. Social status was not linked to infection risk, even after the hurricane. Overall, our study highlights the buffering against infection that affiliative partners may provide, suggesting individuals can compensate for the health costs of sociality by having partners who tolerate their presence.

## Introduction

Parasites and infectious diseases are one of the major costs associated with group living. High population densities and high rates of social interactions can increase parasite transmission, particularly of parasites that rely on social contact between hosts [1]. However, not all individuals in a group are equally likely to get infected, possibly due to differentiated patterns of interactions (*i.e*., not all group members interact with all others) that can modulate levels of exposure and susceptibility to infection [2]. For instance, it is now well established that social partners are not only a potential source of infection but can also be a valuable form of social capital that can positively impact their partner’s health and fitness [3, 4]. Recognising how group-living animals manage the significant trade-off between the cost and benefits of social interactions can bring us closer to understanding the evolution of sociality.

Affiliative social behaviours can have opposing effects on infection risk that may depend on the type of interaction. Grooming - a common affiliative behaviour observed in mammals - involves direct social contact and can increase the likelihood of infection by directly transmitted parasites. For example, in Japanese macaques (*Macaca fuscata*)[5], spider monkeys *Ateles hybridus*) [6], vervet monkeys (*Chlorocebus pygerythrus*) [7] and savannah baboons (*Papio cynocephalus*) [8] individuals with more grooming partners or that engaged more often in grooming interactions were more likely to be infected with nematodes. However, grooming has also been linked to health benefits by directly removing ectoparasites [9, 10]. Grooming is also considered a key behaviour by which social animals establish relationships [11, 12, 13, 14], which can themselves provide indirect health benefits by increasing access to food or shelter [15], or by preventing injuries [16], which can, in turn, promote immunity and resistance to parasites [17, 18]. Similarly, close spatial proximity of individuals might be more favourable for the transmission of parasites, especially those that can survive for extended periods in the environment. For instance, belonging to the same social group and sharing space predicted bacterial and protozoan infection in Verreaux’s sifakas *Propithecus verreauxi*)[19] and Grant’s gazelles (*Nanger granti*)[20], respectively. But spatial proximity is also commonly considered an affiliative interaction reflecting social tolerance, thus, it could alternatively reduce the susceptibility to infection in cases where it provides animals with more opportunities to access the resources necessary to maintain optimal health or reduce exposure by providing access to resources of better quality (*i.e*., uncontaminated food) [21, 22, 19, 23]. All these examples highlight the disease-linked trade-offs that an animal must balance when interacting with conspecifics. Yet, the type of interaction is not the only factor influencing an individual’s exposure to and ability to cope with communicable diseases– the quality of their relationships with others may play a key additional role.

A growing body of research has highlighted the putative importance of weak and strong relationships on an individual’s fitness [24, 25, 26] with potential implications on parasite transmission. Having many social partners with whom infrequent interactions occur (*i.e*., weak relationships) may pose a higher risk of transmission by increasing the diversity of hosts with whom the animal interacts [27]. Strong relationships, where stable social partners frequently interact in an affiliative manner, may also increase the risk of parasite transmission because of longer exposure time [28]. Depending on the context, individuals may prioritise specific relationship types to compensate for the costs associated with social transmission of parasites. For example, weak affiliative relationships may be beneficial in adverse environmental conditions and/or when resources are limited. In macaque species, having more social partners has been associated with enhanced social thermoregulation [29] and heat-stress avoidance [30] when facing harsh winters or a major hurricane, respectively. Strong affiliative relationships can also promote health when expensive returns are required. In vampire bats (*Desmodus rotundus*), for instance, individuals were more likely to donate a blood meal to bats with whom they groomed more frequently [11]. Therefore, both strong and weak affiliative relationships could potentially compensate for the health costs of parasite transmission associated with sociality, but their relative importance may differ depending on environmental conditions.

Another important component of sociality that might also have competing effects on infection risk are dominance hierarchies. On one hand, the way social status (*i.e*., dominance rank) is acquired seems to influence parasitic infection in some mammals. For instance, in rhesus macaques (*Macaca mulatta*) individuals that aggressed others more often had a higher risk of bacterial infection [17]. Similarly, in meerkats (*Suricata suricatta*) animals that were more often victims of aggression were at greater risk of contracting tuberculosis [31]. On the other hand, social status may determine inequalities in access to resources, with individuals higher in the hierarchy at an advantage compared to low-status individuals [32]. Having priority or better access to resources might translate into reduced chances of parasitic infection due to disparities in health status and in the susceptibility to infection [33, 34] or, conversely, into higher chances of infections if it is associated with greater exposure [35]. For instance, a meta-analysis of male and female vertebrates found that high-social status was associated with higher parasite risk [36]. Animals at the highest risk were males from species in which social status determines mating effort, suggesting that high-status and priority of access to mates can also lead to higher exposure to parasites. Thus, how dominance hierarchies impact parasite risk is likely to vary in a similar way as it does to social relationships - depending on the socio-ecological context and how the inequalities it entails translate into health differences.

Growing evidence has contributed to our understanding of the associations between affiliative relationships, social status, and infectious diseases [37, 38, 2]. Yet results to date are mixed and we are still far from a thorough comprehension of the contexts under which social interactions constitute a risk or act as a buffer against parasite infections. For instance, the ecological context can not only shape the distribution and abundance of parasite species [39] but also the aggregation patterns of individuals [30], both with potential consequences on infection dynamics [40]. Further efforts to disentangle some of the factors that may influence the link between sociality and infection including the types of social interactions involved in parasite transmission, the role of different affiliative relationships in buffering infection risk along with individual differences in susceptibility due to age, sex and social status are therefore required. The sociality/infection trade-offs must be particularly relevant in the context of extreme environmental events, which may cause dramatic changes in the environment and in the dynamics of parasite transmission [41].

Here, we studied how social and ecological variation predicted gastro-intestinal parasite burden in a free-ranging population of rhesus macaques. Monkeys in this population self-organize into groups and interact spontaneously with each other, allowing us to explore the consequences of natural variation in social behaviour on infection risk. A natural disaster Hurricane Maria-hit this population in 2017 causing dramatic changes in the environment including a massive decline in vegetation (63%) [30], which provided a unique opportunity to explore how changes in the ecological conditions shape parasite dynamics in a social context. First, to have a better understanding of how changes in the environment affected an individual’s infection risk, we explored 1) whether the hurricane was associated with the risk and intensity of parasite infection. Next, 2) we explored if the risk and the intensity of infection were modulated by social status in general and in the context of the hurricane. Given that social status might determine access to better or cleaner resources [32], especially under the context of the hurricane, we expected that high-status individuals would be less likely to be parasitized. Finally, to disentangle the roles of the type of affiliative social interaction (*i.e*., grooming and proximity) and the quality of relationships (*i.e*., weak and strong) on parasite risk, we tested 3) whether the number of weak grooming and proximity partners and 4) the frequency of interaction with strong grooming and proximity partners predicted infection risk. Given the complexity of the trade-offs detailed above, we did not have clear predictions for these analyses. However, in a general sense, we expected that individuals with more social capital-either in the form of weak partners or strong relationships-would be less susceptible to infection, while the type of interaction involved might determine the types of parasites found.

## Methods

### Subjects and study site

We studied free-ranging rhesus macaques (*Macaca mulatta*) living in the Cayo Santiago field station, Puerto Rico. Animals in this population are provisioned daily with commercial monkey chow and browse on natural vegetation, while water is supplied *ad-libitum* from rainwater collection troughs and is also available from rain water puddles that accumulate naturally. Given that the mean annual population growth rates of the Cayo Santiago macaques are higher than those of wild rhesus populations, live capture and removal of individuals has been implemented by colony management since 1956 [42]. The island is predator-free and there is no regular medical intervention. We studied 67 females and 34 males between the ages of 4 and 26 years (mean = 10.7 years). Macaques belonged to three social groups, where each group represents a single year of data: HH was sampled in 2016 (group size: 95 adults and 13 subadults), KK was sampled in 2018 (group size: 124 adults) and V was sampled in 2020 (group size: 90 adults). Groups KK and V were sampled after they experienced Hurricane Maria.

### Behavioural data collection

Behavioural data were collected using three protocols [43]: 5-min focal animal sample for group HH, group-wide scan sampling for group KK and event sampling for group V. All data collection were done by two experienced observers. Group HH was sampled from August to October 2016 using a previously established protocol for focal sampling [44] that allowed us to have detailed information on social interactions. Specifically, we recorded state behaviours (*i.e*., resting, feeding, travelling) along with grooming, proximity (*i.e*., within 2 m of focal animal) and agonistic interactions. For each of these records, the identities of the focal animal and social partner(s), and the specific behaviour were registered. Agonistic interactions included threat and submissive behaviours along with contact and non-contact aggression. We collected 1.46 ± 0.08 hours of behavioural data per individual in HH. Group KK was sampled from January to October 2018 using scan sampling due to constraints following the animals on the island because of the damage caused by Hurricane Maria, which made landfall in Puerto Rico in September 2017. For this group, we recorded state behaviours, affiliative (i.e., grooming and proximity) and agonistic interactions between all visible adults at 15 min intervals. We collected 538.1 ± 161.3 behavioural events per animal in KK. Group V was sampled from January to December 2020 using event sampling because of restrictions on access to the field site during the COVID-19 pandemic as researchers were allowed in the field for half-days and only every 2-3 days. Specifically, we only recorded information on agonistic encounters, focusing on all the aggressive interactions described above so that we could resolve the dominance hierarchy. We collected 7.13 ± 4.6 agonistic events per individual in V.

### Parasite data collection and identification

Faecal samples (n = 1 per individual) were collected opportunistically in the field from animals living in group V (*n* = 30 samples) and in the laboratory from animals belonging to groups KK (*n* = 16 samples) and HH (*n* = 54 samples), both of which were removed from the population. See Hernandez-Pacheco et al. [42] for details on population control implemented by the field station and Pavez-Fox et al. [18] for details on the removal of individuals used in this study. Although entire groups were scheduled for removal, we could not collect faecal samples from all these individuals. Samples from V were collected a few minutes after defecation within the period of behavioural data acquisition, while samples from HH and KK were collected during necropsy. Each faecal sample was homogenized and stored at room temperature in 10% formalin. In total, we collected 100 samples from 100 individuals: 54 samples collected before the hurricane (34 females, 20 males) and 46 samples collected after the hurricane (32 females, 14 males).

We used the Formalin ethyl-acetate sedimentation technique to extract faecal parasites [45, 46]. We estimated the number of parasites using a wet mount procedure; two drops of a processed sample were placed on a microscopic slide, stained with 5% iodine solution and examined at 10× and 40× magnification. Thick samples were diluted on the slide by adding one drop of 0.9% sodium chloride before the stain. We examined two drops of processed sample twice per sample, thus each individual had two records. For one male, we did not have enough sample to perform the procedure twice, so only one was included. Parasite taxa were identified by direct observation based on their morphology (*e.g*., shape, colour, size) [47, 48]. Larvae were rarely seen, and we, therefore, identified the presence of helminths based on the morphology of eggs. We identified five parasite taxa across all samples including one protozoa (*Balantidium coli*) and four nematodes (*Ascaris lumbricoides, Ancyclostoma* spp., *Strongyloides fuelleborni, Trichuris trichiura*). For our analyses, we focused on the three most prevalent parasites (Fig. 1; *B. coli, T. trichiura* and *S. fuelleborni*) (details on prevalence are presented in the Results), all of which detected in previous studies in this population [49, 50]. We estimated two measures of infection per host: the presence of infection per parasite taxa (*i.e*., infection risk: 0 = absent, 1 = present) and intensity of infection (*i.e*., count of parasites on infected hosts) for nematodes and protozoa separately, based on differences on transmission routes [51].

**Figure 1.**
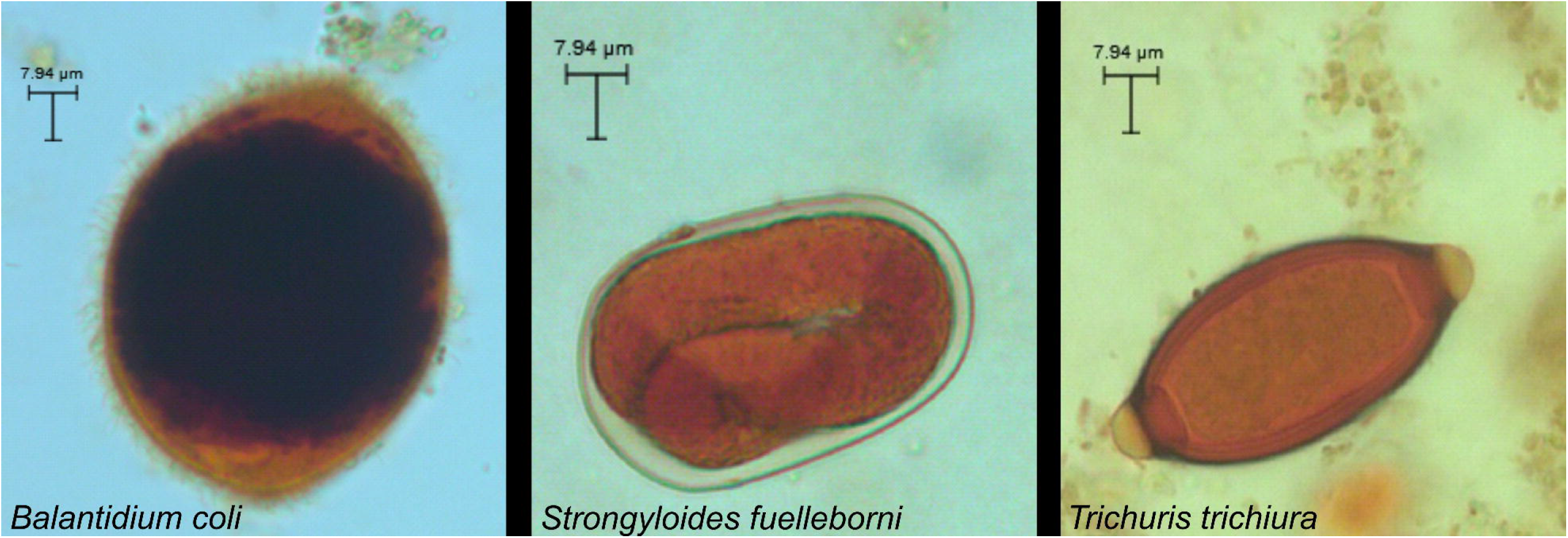
Most prevalent parasite species found in the faecal samples of the Cayo Santiago macaques, including a protozoan (*B. coli* trophozoite) and two nematodes (*S. fuelleborni* and *T. trichiura* eggs). Photos taken with a light microscope camera.

### Dominance hierarchies

To determine an individual’s social status we computed dominance hierarchies by group and separately for males and females [52, 53, 54]. Our approach is based on the fact that, in this species, males and females acquire social status differently. Females are philopatric and form maternally inherited stable linear dominance hierarchies, where daughters acquire rank just below their mothers [55]. In contrast, males typically disperse from the natal group and acquire rank in the new group by physical contest and tenure [56]. We built independent hierarchies for the three social groups, using the outcomes of win-loss agonistic encounters from focal/scan sampling and *ad-libitum* observations, with known maternal relatedness used to resolve behavioural gaps in the female hierarchy. To account for variation in group sizes, dominance rank was defined as the percentage of group mates from a subject’s sex that they outranked, where 100% corresponded to the highest-ranking animal [57].

### Social networks

Using proximity and grooming interactions, we constructed social networks for groups that had data on affiliative interactions (*n* = 2 groups). We included all non-juveniles for which we had data: all adult animals from group KK and adults plus subadults for group HH. We built separate networks for each interaction type. We focused on two network metrics that allowed us to delineate relationship types: the number of weak connections and the frequency of interaction (*i.e*., their relationship strength) with strong partners. Weak connections were quantified as the number of social partners with whom an individual engaged in infrequent affiliative interactions, while the frequency of interaction with strong partners quantifies the time invested in strong relationships [24, 25]. The thresholds used to establish weak and strong partnerships are explained below.

We generated undirected weighted Bayesian networks using the BISoN framework and bisonR package [58]. This framework allowed us to account for uncertainty in the edges connecting individuals in the network based on how often they were sampled and, more importantly, propagate this uncertainty to subsequent analyses. In all our networks we modelled the uncertainty around the edges using as prior a beta distribution with alpha = 0.1 and beta = 1. For the proximity networks, an edge represented the number of times a pair of individuals were observed in proximity relative to the total observation effort for the dyad (*i.e*., total scans individual A + total scans individual B). For the KK grooming network, an edge between individuals represented the number of scan records a dyad engaged in grooming interactions relative to their total observation effort (*i.e*., total scan records individual A + total scan records individual B). For the HH grooming network, edges represented the time a pair of individuals engaged in grooming interactions relative to the total time that dyad was observed. Given that there is no natural statistical model for duration data [58], the time spent grooming and the sampling effort for a dyad were converted to counts by dividing each of these terms by the length of a focal period (5-mins) to make sure each count represented independent sampling events.

Networks generated with BISoN include edges between all dyads by default, as it assumes non-zero probability for all potential interactions, even if that probability is exceedingly small. To compute the number of weak partners, we therefore defined a threshold that allowed us to differentiate dyads that did interact versus those that did not based on the minimum observed edge weight in each network. That is, for each of the posterior samples, dyads with a BISoN edge weight above or equal to the minimum empirical edge weight were kept and those below that value were excluded from the computation of network metrics. Strong partners were defined as dyads that had an edge weight within the upper quantile (*i.e*., 75% and above) of all the existent connections in a network, while weak partners were those dyads that had edge weights values below this quantile (see Fig. S1 for visualisation of both thresholds on each network). An individual’s number of weak partners was the number of edges they had that were classed as ‘weak’. An individual’s relationship strength to strong partners was computed by summing the weights of their edges that were classed as ‘strong’ connections. All network metrics were set to range between 0 and 10 by dividing by the maximum value of that metric for the group and multiplying it by 10. By doing so, we accounted for possible group differences attributed to sampling methods because the network metrics were scaled relative to other individuals within a group.

### Statistical analyses

All statistical analyses were carried out in R v4.3 using the *brms* package for Bayesian statistics [59]. For all models of infection risk, the dependent variables were the binary presence (1) or absence (0) of a given parasite species in the sample. For all models of the intensity of infection, the dependent variables were the count of parasites per taxa (protozoa and nematodes). To quantify the intensity of infection only infected animals were included [51] (*n* unique infected individuals: protozoa = 60, nematodes = 35), thus we truncated at zero the dependent variable in our models. Additionally, for the intensity of protozoa, we right-censored the dependent variable, as we only quantified up to 60 parasites per sample even in cases where animals had more parasites (Out of 199 records from 100 samples, 9 were censored).

### Risk and intensity of infection before and after hurricane

To test whether parasite infection risk differed before and after the hurricane, we used linear mixed models with a binomial distribution (‘Bernoulli’ in *brms* environment), running one model per parasite species. The dependent variable was the presence or absence of a given parasite taxon. As predictors, we included hurricane status, where 0 = sampled before and 1 = sampled after the hurricane, along with the age and sex of the animal. We also included a fixed effect for the season when the sample was collected (rainy vs dry season) to account for changes in precipitation and temperature that might influence parasite dynamics [39].

To determine if the hurricane influenced the intensity of parasite infection, we used linear mixed models with a Poisson distribution. We included in the model hurricane status as the main predictor, along with sex, age, and season as covariates. We tested for interaction terms between hurricane status and all the other fixed terms and retained them when evidence of an effect was detected. For all models of infection risk and intensity of infection, we included a random effect for ID to account for repeated records per individual.

### Effect of social status on the risk and intensity of infection

To determine if social status influenced the infection risk overall and in the context of the hurricane, we used logistic models where the dependent variable was the presence/absence of infection per parasite species. As fixed effects, we included social status, hurricane status (0 = pre, 1 = post), age, and sex. We accounted for repeated records per individual by including a random effect for animal ID. To test if social status influenced the intensity of infection, we modelled our dependent variables as described above. Fixed effects and the random effect followed the same format as the models for infection risk. We first ran a set of models to test if social status buffered the impact of the hurricane on parasitic infections (interaction among those predictors), and then we ran a second set of models where no interactions between predictors were included to establish the magnitude of main effects.

### Effect of weak and strong relationships on infection risk

For all the analyses that included social network metrics, animals from group V were excluded, as we did not have behavioural observations on affiliative interactions for this group. This resulted in a smaller sample size (*n* = 70), especially post-hurricane (*n* = 16), that restricted our ability to test the impact of Hurricane Maria in relation to affiliative relationships.

**Table 1:** Dependent variables, fixed and random effects used in the models.

| Variable | Type | Description |
| --- | --- | --- |
| *B. coli* infection | Dependent | Presence or absence of *B. coli* infection |
| *S. fuelleborni* infection | Dependent | Presence or absence of *S. fuelleborni* infection |
| *T. trichiura* infection | Dependent | Presence or absence of *T. trichiura* infection |
| Intensity of protozoa infection | Dependent | Count of *B. coli* parasites in infected host |
| Intensity of nematode infection | Dependent | Count of *S. fuelleborni* and *T. trichiura* parasites in infected host |
| Social status | Fixed effect | Percentage of same-sex individuals outranked in the group |
| Number of weak partners | Fixed effect | Number of proximity or grooming partners with whom the focal interacted infrequently* |
| Strength to strong partners | Fixed effect | Rate of engagement in proximity or grooming interactions with strong* social partners |
| Hurricane status | Fixed effect | Sampled before or after the hurricane Maria |
| Age | Fixed effect | Age of the individual when it was sampled |
| Sex | Fixed effect | Sex of the animal |
| Season | Fixed effect | Sample collected during the wet or the dry season |
| Animal ID | Random effect | Identification of the macaque from which the sample was taken |

\*relative to the upper quantile of the group, where infrequent represents a rate of interaction (i.e., edge weight) below the 75% quantile and strong refers to a rate above or equal to it. Prevalence was modelled as binary (0/1). The count for intensity excluded zeroes. Hurricane status was modelled as binary (0/1). All continuous predictors were z-scored.

The exclusion of group V also resulted in a substantial reduction in the number of infected individuals (*n* unique infected individuals: protozoa = 36, nematodes = 21), therefore we did not test the effect of affiliative relationships on the intensity of infection or the effect of interactions between our predictors. All the samples for the remaining groups (HH and KK) were collected during the same period (between October and November in their respective years) so season was not included in these models.

To test if individuals with a greater number of weak relationships had a higher risk of parasite infection, we used logistic models where the dependent variable was the presence/absence of infection per parasite species. We tested the effect of the number of weak partners on parasite presence for grooming and proximity in separate models to avoid over-parametrization with our limited sample size. In all the models, our main predictor was the number of weak partners with covariates for social status, age, and sex.

To test if individuals with stronger social relationships had a reduced risk of parasite infection, we used logistic models where the dependent variable was the presence/absence of infection per parasite species. As main predictors, we included the strength of relationships to strong partners (separate models for grooming and proximity) along with an individual’s social status, age, and sex as covariates. We accounted for repeated records per individual by including a random effect for animal ID.

### Bayesian model specifications

In all the models we used weakly informative priors, which are recommended over flat priors to avoid overfitting issues when sample sizes are small and no prior knowledge of the relationship between the dependent variable and predictors is assumed [60]. Specifically, we used a t-student distribution with a mean of 0, 5 degrees of freedom and a standard deviation of 2.5 for all our fixed effects. We opted for a t-student distribution as it is less sensitive to outliers or skewed data compared to a normal distribution. Using weakly informative priors that assign more weight to the absence of an effect (mean = 0) also helps to mitigate the need to account for multiple testing when repeated tests of the same dataset are performed [61], like in our case. We z-scored all the continuous predictors to improve sampling efficiency and to match prior specifications for the intercept (mean-centred at 0). We assessed model convergence by examining the R-hat values (⍰ 1), effective sample sizes (> 1000) and visual inspection of the chains. We checked the goodness of fit of the models by using the pp_check function from the brms package, which allowed us to do posterior predictive checks by comparing the data from the posterior distribution of the models with the observed data. In the case we detected an interactive effect of our predictors, we used the emmeans R package [62] to perform a post-hoc test. We reported means as point estimates, standard error (SE) and 89% credible intervals of the posterior distribution. Evidence for an effect was determined based on the degree of overlap between the credible interval and zero (*i.e*., 89% non-overlapping reflecting strong evidence for an effect). For post-hoc tests, we reported the median and the 89% highest posterior density interval (HPD). All the parameters included in the models can be found in Table 1 and model specifications in Table S1.

## Results

The most common parasites detected in our samples was a protozoa (*Balantidium coli*), which was present in 60 of the animals sampled (60% prevalence) and two nematode species: *Trichuris trichiura* (24% prevalence) and *Strongyloides fuelleborni* (23% prevalence). We also identified two other helminth taxa (*Ascaris lumbricoides, Ancyclostoma* spp.), but these were rarely seen and thus not included in downstream analyses. Twenty-five out of the 100 individuals sampled did not harbour any parasites.

### Infection risk before and after hurricane

*B. coli* was found in 38 individuals before the hurricane (70% prevalence) compared to 22 animals post-hurricane (48% prevalence). *S. fuelleborni* was present in 8 animals before the hurricane (15% prevalence) and 15 individuals after (33% prevalence), while *T. trichiura* had a prevalence of 15% before the hurricane (present in 8 animals) and of 35% post-hurricane (present in 16 individuals).

Overall, the hurricane significantly impacted the risk of infection but this varied with the parasite species examined and with the age and sex of the macaque host. We found that *B. coli* infection was associated with the hurricane independent of the sex of the host and the season but dependent on the host’s age (Fig. 2A; Log-Odds age*post-hurricane = −0.72, SE = 0.27, 89%CI = −1.23, −0.33; Table S2). The risk of infection with *B. coli* had a positive relationship with an individual’s age before the hurricane (post hoc test: Log Odds pre-hurricane = 0.44, 89% HPD = 0.0784, 0.842) but was not linked to age after the hurricane (post hoc test: Log Odds post-hurricane = −0.274, 89% HPD = −0.689, 0.106). Individuals sampled after the hurricane had less intense *B. coli* infections compared to animals sampled before the hurricane (Log-Odds post-hurricane = −0.91, SE = 0.49, 89%CI = −1.71, −0.12; Table S3).

**Figure 2.**
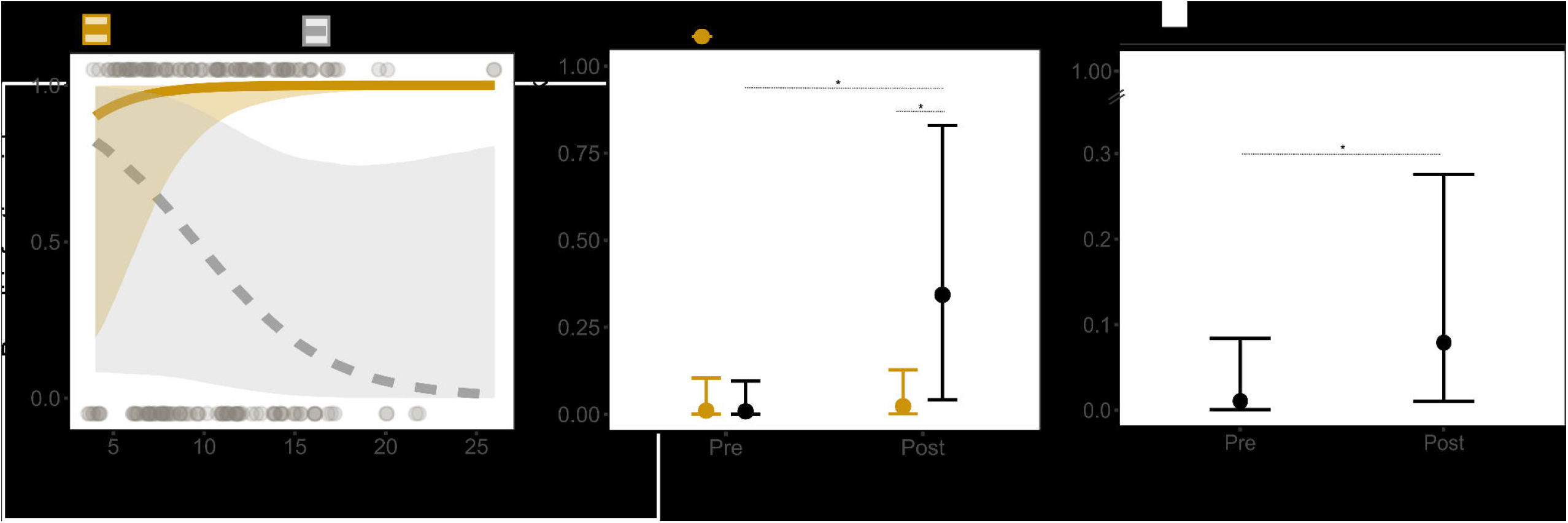
Infection risk before and after the hurricane. **A)** Prevalence of *B. coli* as a function of an individual’s age and hurricane status (pre-hurricane: grey, post-hurricane: brown). Shaded area represents 89% credible interval and lines the median. Grey points reflect the raw data, where those on the top indicate the presence of infection and those on the bottom, the absence of it. **B)** Prevalence of *S. fuelleborni* as a function of sex and hurricane status. **C)** Prevalence of *T. trichiura* before and after the hurricane. Errors bars represent the 89% credible interval and point estimates, the medians. Evidence for an effect is indicated with an asterisk.

The hurricane was also associated with the prevalence of *S. fuelleborni*. This was irrespective of individual age and season but in a sex-specific manner (Fig. 2B; Log-Odds sexM*post-hurricane = 3.3, SE = 1.88, 89% CI = 0.68, 6.73; Table S4). Only male’s infection risk changed with the impact of the hurricane. Males were more likely to be infected with *S. fuelleborni* after than before the hurricane (post hoc test: Males pre vs post-hurricane Log-Odds = −4.04, 89% HPD = −7.16, −1.4), while females had a similar risk of infection before and after hurricane Maria (post-hoc test: Females pre vs post-hurricane Log-Odds = −0.71, 89% HPD = −2.83, 1.07). After the hurricane the risk of infection with *T. trichiura* was higher irrespective of the age, season, and sex of the animal (Fig. 2C; Log-Odds post-hurricane = 2.06, SE = 1.04, 89%CI = 0.47, 3.93; Table S5), but it did not affect the intensity of nematode infection (Log-Odds post-hurricane = 0.46, SE = 0.44, 89%CI = −0.27, 1.19; Table S6).

### Effect of social status on risk and intensity of infection

We did not find evidence for a buffering effect of social status on parasite infection overall or in the context of the hurricane. Social status was not associated with infection risk (*B. coli*: Log-Odds status = −0.18, SE = 0.73, 89%CI = −1.38, 1.03, Table S7; *S. fuelleborni*: Log-Odds status = 0.33, SE = 0.6, 89%CI = −0.65, 1.38, Table S9; *T. trichiura*: Log-Odds status = − 0.22, SE = 0.52, 89%CI = −1.12, 0.62, Table S10) or with the intensity of infection overall (*B. coli*: Log-Odds status = 0.15, SE = 0.19, 89%CI = −0.17, 0.46, Table S8; nematodes: Log-Odds status = 0.03, SE = 0.23, 89%CI = −0.35, 0.4, Table S11). We also did not find evidence for a modulatory effect of the hurricane on the association between social status and infection risk (*B. coli*: Log-Odds status*hurricane = −1.55, SE = 1.32, 89%CI = −3.83, 0.55, Table S11; *S. fuelleborni*: Log-Odds status*hurricane = 0.48, SE = 1.12, 89%CI = −1.32, 2.36, Table S13; *T. trichiura*: Log-Odds status*hurricane = −0.3, SE = 0.95, 89%CI = −1.9, 1.26, Table S14) nor on the intensity of infection (*B. coli*: Log-Odds status*hurricane = 0.24, SE = 0.39, 89%CI = − 0.39, 0.86, Table S12; nematodes: Log-Odds status*hurricane = −0.02, SE = 0.52, 89%CI = − 0.87, 0.81, Table S15).

### Effect of weak relationships on infection risk

The number of weak social partners in the proximity network was negatively associated with the prevalence of *B. coli*. Macaques that had a greater number of weak proximity partners were less likely to be infected than individuals with fewer weak partners (Fig. 3A; Log-Odds = −2.03, SE = 1.118, 89%CI = −4.16, −0.18; Table S17). No effect of the number of weak partners in the grooming network on *B. coli* infection risk was detected (Log-Odds = −1.71, SE = 1.39, 89%CI = −4.19, 0.46; Table S16).

**Figure 3.**
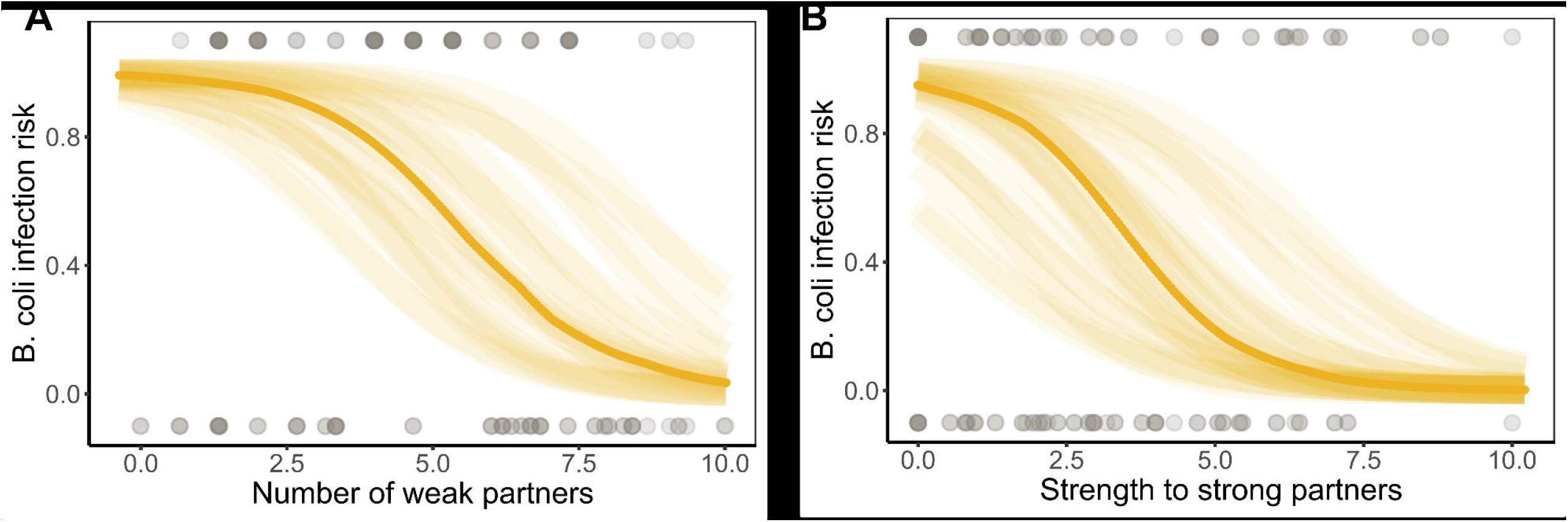
Effect of **(A)** the number of weak partners and **(B)** the strength of relationship to strong partners on *B. coli* infection prevalence. In both plots the solid yellow line represents the median prevalence in samples and the shaded area corresponds to values from 20 draws from the posterior distribution within the 89% credible interval. Raw data are depicted with grey points (top: infected, bottom: non-infected).

When we tested the effect of the number of weak partners on infection risk with *S. fuelleborni* and *T. trichiura* the estimates were nearly identical between both parasite species (Tables S18-S21), which seemed to indicate problems of over-parametrization, especially with the lower prevalence for these infections. Therefore, we re-ran these models only including the number of weak partners as predictors (*i.e*., univariate models), as all other covariates did not have an effect on prevalence when themselves tested in univariate models (Tables S22-S27). We found that infection risk with *S. fuelleborni* was not predicted by the number of weak partners in the proximity or grooming networks (grooming: Log-Odds = 0.26, SE = 0.73, 89%CI = −0.94, 1.52, Table S28; proximity: Log-Odds = 0.48, SE = 0.66, 89%CI = −0.56, 1.67, Table S29). Similarly, the number of weak partners did not predict the prevalence of *T. trichiura* in the grooming or proximity networks (grooming: Log-Odds = 0.04, SE = 0.64, 89%CI = −1.08, 1.11, Table S30; proximity: Log-Odds = 0.37, SE = 0.56, 89%CI = −0.56, 1.35, Table S31). In other words, having more weak proximity partners was associated with a reduced risk of protozoan infection but had no relationship with the likelihood of infection with nematodes.

### Effect of strong relationships on infection risk

Macaques that were more often observed in proximity to their strong partners were less likely to be infected with *B. coli* (Fig. 3B; Log-Odds = −2.07, SE = 1.29, 89% CI = −4.36, −0.02; Table S33). No effect of strong relationships with grooming partners on the infection risk with this protozoan was detected (Log-Odds = −1.06, SE = 1.41, 89% CI = −3.47, 1.24; Table S32).

The models for nematode infection risk showed signs of over-parametrization (Tables S34-S37), so as mentioned above, we re-ran univariate models including only the network metrics as predictors. We found that the frequency of observations spent in proximity or grooming with strong partners did not influence the risk of infection with *S. fuelleborni* (grooming: Log-Odds = 0.01, SE = 0.79, 89% CI = −1.4, 1.32, Table S38; proximity: Log-Odds = 0.23, SE = 0.64, 89% CI = −0.84, 1.36, Table S39) or *T. trichiura* (grooming: Log-Odds = 0.2, SE = 64, 89% CI = −0.87, 1.29, Table S40; proximity: Log-Odds = 0.25, SE = 0.56, 89% CI = −0.72, 1.19, Table S41). Overall, we only found evidence that strong relationships in the proximity network predicted infection risk, showing that animals that shared space with their strong partners more often were less likely to be infected with a protozoan.

## Discussion

Our results provide evidence for a buffering effect of affiliative relationships on infection risk. We found that social status did not play a role in mitigating infection overall or after the impact of a major hurricane, and that grooming relationships, whether weak or strong, were not associated with parasite infection. But those individuals that spent more time with weak and strong social partners were less likely to be infected with a protozoan. Together, these results suggest that social tolerance is one way by which affiliative partners can help prevent infection from environmental parasites.

We found that environmental changes caused by a major disaster are not always associated with a greater risk of parasite transmission. Our results instead suggest that the relationship between parasite transmission and environmental upheaval depends on the life cycle of the parasite under study. For example, we found that the risk and the intensity of infection with the protozoa *B. coli* in the Cayo Santiago population were higher before Hurricane Maria compared to after it. Optimal environmental conditions for the infective stage of *B. coli* (*i.e*., cysts) are humid areas protected from direct sunlight [63], which were very scarce after the hurricane given the massive loss of vegetative cover [30, 64]. Before the hurricane, when *B. coli* was more prevalent, the likelihood of infection was higher for older individuals compared to younger ones, which could reflect increased susceptibility due to immunosenescence [65] or also be the consequence of higher exposure if older individuals are less able to access uncontaminated resources.

Infection risk with nematodes was also associated with the hurricane. Males were more likely to be infected with *S. fuelleborni* than females after the hurricane. Given that previous evidence in this population found that Hurricane Maria was not associated with sex differences in immune-gene expression [64], changes in the environment are more likely to have led to higher parasite exposure in males, instead of this result being the consequence of exacerbated immunosuppressive effects of testosterone after a natural disaster [66, 67]. Infection risk with *T. trichiura* was also higher after Hurricane Maria. As the transmission of both nematode species can occur by animals touching infected areas on the soil and then ingesting the parasite eggs [68], enhanced exposure after the hurricane could be due to macaques being more clustered in shade and engaging more in social behaviour [30]. Nevertheless, these results should be interpreted with caution, as the prevalence of nematodes was relatively low, leading to greater uncertainty in our models. Compared to a previous study of in this same population [49], we found a similar prevalence of *T. trichiura* but lower *S. fuelleborni*, which could be attributed to group differences and/or the smaller sample size of our study (*n* = 256 vs 100 in our study). Differences in the faecal collection method could also explain a lower prevalence of *S. fuelleborni* as most of our samples were collected during necropsy and therefore were less likely to be contaminated with eggs from these nematodes that were already on the soil. Although we would expect collection, method to impact results for both types of nematodes in the same way, which does not appear to be the case.

Contrary to our predictions, we did not find a buffering effect of social status on infection risk, either in general or in the aftermath of a hurricane. These results contrast growing evidence that health differences are associated with inequalities linked to social status [33, 3]. In several vertebrate species, high-status individuals, especially males, have a higher parasitic load from contact and environmentally transmitted parasites, which has been posited to be due to greater exposure given their priority of access to resources [35, 36]. That is, dominant individuals are usually the ones that take most of the food increasing their risk of infection with parasites transmitted via the faecal-oral route [69], and/or have more mating opportunities, which increases the chances of contact transmission of parasites [70]. Yet, our results are in line with those from a previous study in this population that showed that immune function was not influenced by social status [18]. The macaques on Cayo Santiago are food provisioned, thus the chances of eating contaminated food might be lower than in wild populations and social status does not strongly determine reproductive skew [71]. Together, these results suggest that social status is not a strong determinant of susceptibility and exposure to parasites in these animals, even under adverse ecological conditions.

Proximity interactions were associated with infection risk, but grooming was not. We were able to disentangle potential transmission routes by quantifying different types of social interactions and their effects on infection risk for different parasite species. Sharing space was associated with a reduced likelihood of *B. coli* infection. This parasite is commonly found in contaminated food or water [63]. Monkeys in the Cayo Santiago population are food provisioned with monkey chow and have *ad-libitum* access to rainwater in the form of puddles and, rainwater that runs from passive water collectors into drinking troughs [72]. Yet, competition might prevent some individuals to access these resources [73]. Our results suggest that proximity to other individuals might enable access to better quality or cleaner resources by means of social tolerance [21], which may reduce the exposure to *B. coli*.

We did not find a relationship between infection risk and grooming interactions, contrasting previous evidence from other systems where this behaviour has been associated with higher infection risk [5, 6, 7, 8]. Yet, we cannot disregard that our reduced sample (*e.g*., one sample per animal vs multiple samples) and the low prevalence of nematodes in our animals - which are commonly the parasites associated with social contact transmission - explain these results.

Both weak and strong relationships in the proximity network were associated with a buffering effect against *B. coli* infection. This suggests that both strategies, either having many partners or relying on strong partners allow individuals to avoid exposure to this protozoan. In conditions where feeding resources are not scarce, like in the Cayo Santiago population, it is likely that tolerating social partners around feeding or drinking areas does not translate into major costs for the animals. This might explain why we did not find distinct effects between weak and strong proximity partners, as not only strong partners but also weak ones might provide access. Surprisingly, we did not find a buffering effect of weak or strong grooming relationships on infection risk. Previous evidence in this system relying on a similar dataset has shown that the number of grooming partners - a measure that closely reflects the number of weak grooming connections (Fig. S2) - was associated with reduced inflammation levels [18]. However, our results seem to suggest that this does not come about because of a reduced likelihood of being parasitized.

## Conclusion

The results of our study highlight how affiliative interactions - specifically weak and strong social partners-can buffer infection risk. Although social inequalities are usually thought to stem from differences in social status, our results emphasize that affiliative relationships also constitute a valuable resource that might compensate for some of the costs associated with living in groups. These findings add more evidence to a growing body of research on the means by which social capital can benefit an individual’s health and ultimately survival.

## Ethical statement

This research complied with protocols approved by the Institutional Animal Care and Use Committee (IACUC) of the University of Puerto Rico (protocol no. A6850108) and by the University of Exeter School of Psychology’s Ethics Committee. The CPRC’s Animal Care and Use Program are evaluated and approved by the IACUC. Pain and distress are assessed as part of the program. Every protocol used in research, teaching, testing or as part of the daily management of the Center, is evaluated by the IACUC using USDA pain and distress categories.

## Author contributions

Conceptualization: M.A.P-F, L.J.N.B, C.M.E-R, J.P.H. and N.S-M.; Methodology: M.A.PF, C.M.E-R, J.D.A.H.; Resources: L.J.N.B, J.P.H., N.S-M., M.J.M., M.L.P., A.R-L., M.I.M.; Data curation: M.A.P-F, C.M.E-R, J.E.N-D, D.P.; Writing-Original Draft: M.A.P-F; Writing-Review & Editing: M.A.P-F, L.J.N.B., J.P.H., N.S-M.; Supervision: L.J.N.B.

## Acknowledgments

We thank the Caribbean Primate Research Center (CPRC) for the permission to undertake research on Cayo Santiago, along with many interns who assisted in behavioural data collection. Additionally, we thank members of the Brent lab, CRAB lab and Network Club for their helpful suggestions.

