## Supplemental Material for "Cost and benefits of being social: examining the influence of sociality on faecal parasites in free-ranging rhesus macaques"

### Supplementary Information

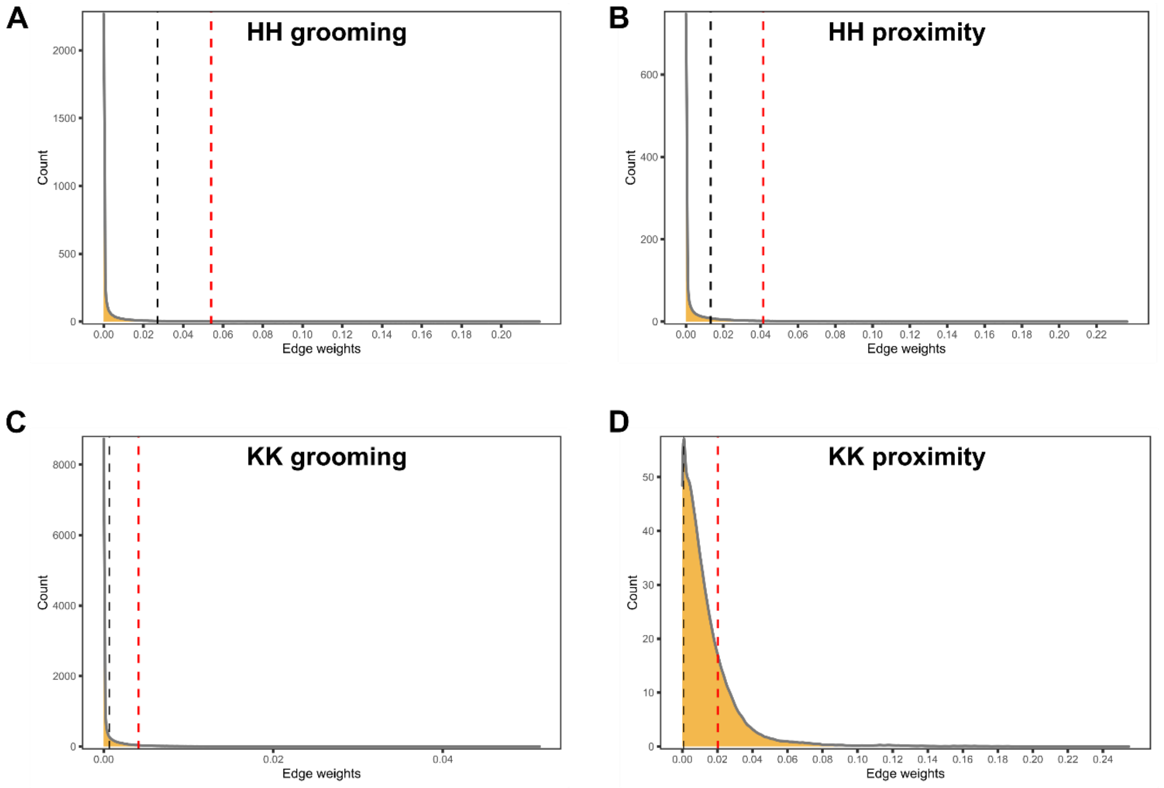


**Figure S1.** Thresholds used to define existent connections (black dotted line) and strong partners (red dotted line) for grooming and proximity networks generated with BISoN.

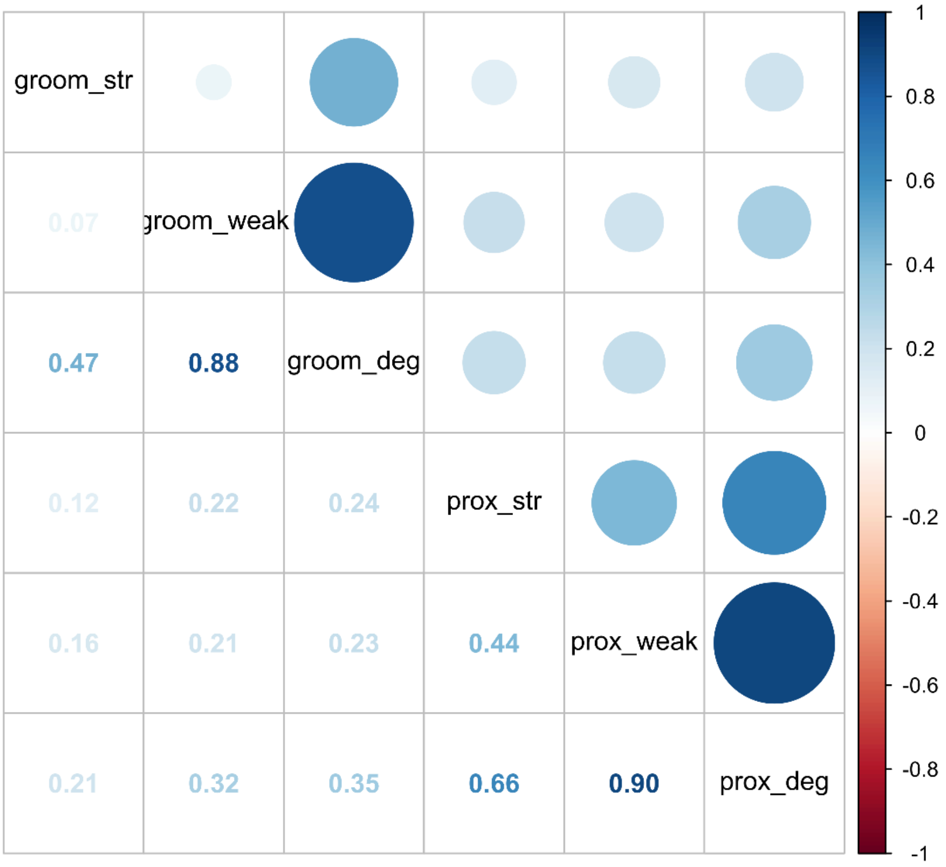


**Figure S2.** Correlation between the networks metrics used in this study obtained from 50 draws from the posterior distribution. We additionally included "degree" to illustrate that the number of weak partners closely resembles this measure of social capital.

**Table S1.** Models specifications.

| Question | Model |
|---|---|
| Did the hurricane affect the prevalence and intensity of parasite infections? | prev_coli ~ hurricane*age + sex + season + (1\|id)<br>prev_strong ~ hurricane*sex + age + season + (1\|id)<br>prev_trichu ~ hurricane + sex + age + season + (1\|id) |
| | int_proto ~ hurricane + sex + age + season + (1\|id)<br>int_nema ~ hurricane + sex + age + season + (1\|id) |
| Does social status influence the prevalence and intensity of parasite infections overall and in the context of the hurricane? * | prev_coli ~ social status + hurricane + sex + age + season + (1\|id)<br>prev_strong ~ social status + hurricane + sex + age + season + (1\|id)<br>prev_trichu ~ social status + hurricane + sex + age + season + (1\|id) |
| | int_proto ~ social status + hurricane + sex + age + season + (1\|id)<br>int_nema ~ social status + hurricane + sex + age + season + (1\|id) |
| Did the number of weak connections an individual had predict infection risk? | prev_coli ~ weak_groom + sex + age + social status + (1\|id)<br>prev_strong ~ weak_groom + sex + age + social status + (1\|id)<br>prev_trichu ~ weak_groom + age + sex + social status + (1\|id) |
| | prev_coli ~ weak_prox + sex + age + social status + (1\|id)<br>prev_strong ~ weak_prox + sex + age + social status + (1\|id)<br>prev_trichu ~ weak_prox + age + sex + social status + (1\|id) |
| Did the frequency of interaction with strong partners predict infection risk? | prev_coli ~ groom_topstr + sex + age + social status + (1\|id)<br>prev_strong ~ groom_topstr + sex + age + social status + (1\|id)<br>prev_trichu ~ groom_topstr + age + sex + social status + (1\|id) |
| | prev_coli ~ prox_topstr + sex + age + social status + (1\|id)<br>prev_strong ~ prox_topstr + sex + age + social status + (1\|id)<br>prev_trichu ~ prox_topstr + age + sex + social status + (1\|id) |

prev_coli = presence/absence of *Balantidium coli,* prev_strong = presence/absence of *Strongyloides fuelleborni*, prev_trichu = presence/absence of *Trichuris trichiura,* hurricane = sampled before or after the hurricane Maria, sex = an indiviual's sex when sampled, age = animal's age when sampled, season = sampled collected during the wet or rainny season, int_prot = intensity of protozoan infection (*B. coli*), id = animal ID, int_nema = intensity of nematode infection (*S. fuelleborni + T. trichiura*), social status = rank of an animal relative to all same sex members of its group, weak_groom = number of weak grooming connections, weak_prox = number of weak proximity connections, groom_topstr = strength to strong grooming partners, prox_topstr = strength to strong proximity partners.

*To test for an effect of social status in the context of the hurricane models were ran with an interaction between these predictors.

**Table S2.** Prevalence of *B. coli* before and after the hurricane as a function of age.

| | Infection risk | | |
|---|---|---|---|
| *Predictors* | *Log-Odds* | *std. Error* | *CI (89%)* |
| Intercept | 0.46 | 2.84 | -4.31 – 5.21 |
| age | 0.44 | 0.23 | 0.10 – 0.88 |
| post_hurricane: post_hurricane1 | 1.98 | 2.50 | -1.66 – 6.65 |
| sex: M | 0.43 | 1.31 | -1.76 – 2.61 |
| season: rainy | -2.44 | 1.80 | -5.67 – 0.28 |
| age:post_hurricane1 | -0.72 | 0.27 | -1.23 – -0.33 |
| $N_{id}$ | 100 | | |
| Observations | 199 | | |

scaleage = z-standardized age; post_hurricane1 = after the hurricane; sex:M = males; season:rainy = wet season.
Model included a random effect for animal id.

**Table S3.** Intensity of *B. coli* infection before and after the hurricane.

| | Infection risk | | |
|---|---|---|---|
| *Predictors* | *Log-Mean* | *std. Error* | *CI (89%)* |
| Intercept | 1.98 | 0.63 | 0.97 – 2.98 |
| post_hurricane: post_hurricane1 | -0.91 | 0.49 | -1.70 – -0.12 |
| season: rainy | 0.05 | 0.59 | -0.90 – 0.98 |
| scaleage | 0.26 | 0.17 | -0.02 – 0.55 |
| sex: M | 0.28 | 0.37 | -0.33 – 0.88 |
| $N_{id}$ | 60 | | |
| Observations | 107 | | |

scaleage = z-standardized age; post_hurricane1 = after the hurricane; sex:M = males; season:rainy = wet season.
Model included a random effect for animal id. Analysis only included animals with count above zero (zero truncated) and was right censored (count = 60).

**Table S4.** Prevalence of *S. fuelleborni* infection before and after the hurricane.

| | Infection risk | | |
|---|---|---|---|
| *Predictors* | *Log-Odds* | *std. Error* | *CI (89%)* |
| Intercept | -4.53 | 1.70 | -7.63 – -2.15 |
| post_hurricane: post_hurricane1 | 0.71 | 1.19 | -1.14 – 2.79 |
| sex: M | -0.24 | 1.29 | -2.53 – 1.77 |
| scaleage | -0.41 | 0.57 | -1.36 – 0.49 |
| season: rainy | -0.18 | 1.15 | -2.13 – 1.62 |
| post_hurricane1:sexM | 3.30 | 1.88 | 0.68 – 6.73 |
| N id | 100 | | |
| Observations | 199 | | |

scaleage = z-standardized age; post_hurricane1 = after the hurricane; sex:M = males; season:rainy = wet season. Model included a random effect for animal id.

**Table S5.** Prevalence of *T. trichiura* infection before and after the hurricane.

| | Infection risk | | |
|---|---|---|---|
| *Predictors* | *Log-Odds* | *std. Error* | *CI (89%)* |
| Intercept | -4.54 | 1.53 | -7.49 – -2.39 |
| post_hurricane: post_hurricane1 | 2.06 | 1.04 | 0.47 – 3.93 |
| scaleage | 0.42 | 0.47 | -0.34 – 1.23 |
| sex: M | 0.98 | 0.92 | -0.49 – 2.62 |
| season: rainy | -0.41 | 1.13 | -2.35 – 1.39 |
| N id | 100 | | |
| Observations | 199 | | |

scaleage = z-standardized age; post_hurricane1 = after the hurricane; sex:M = males; season:rainy = wet season. Model included a random effect for animal id.

**Table S6.** Intensity of nematode infection before and after the hurricane.

| | Infection risk | | |
|---|---|---|---|
| *Predictors* | *Log-Mean* | *std. Error* | *CI (89%)* |
| Intercept | 0.69 | 0.59 | -0.29 – 1.64 |
| post_hurricane: post_hurricane1 | 0.46 | 0.44 | -0.27 – 1.19 |
| season: rainy | 0.42 | 0.47 | -0.37 – 1.18 |
| sex: M | -0.25 | 0.39 | -0.87 – 0.38 |
| scaleage | -0.19 | 0.20 | -0.51 – 0.13 |
| $N_{id}$ | 35 | | |
| Observations | 52 | | |

scaleage = z-standardized age; post_hurricane1 = after the hurricane; sex:M = males; season:rainy = wet season. Model included a random effect for animal id. Analysis only included animals with count above zero (zero truncated).

**Table S7.** Prevalence of *B. coli* infection as a function of social status (main effect).

| | Infection risk | | |
|---|---|---|---|
| *Predictors* | *Log-Odds* | *std. Error* | *CI (89%)* |
| Intercept | 3.07 | 1.94 | -0.00 – 6.45 |
| scaleoutrank_perc | -0.18 | 0.73 | -1.38 – 1.03 |
| scaleage | 0.54 | 0.76 | -0.68 – 1.86 |
| sex: M | 0.18 | 1.35 | -2.02 – 2.41 |
| post_hurricane: post_hurricane1 | -3.39 | 1.60 | -6.41 – -0.96 |
| season: rainy | -1.25 | 1.64 | -4.08 – 1.46 |
| $N_{id}$ | 100 | | |
| Observations | 199 | | |

scaleoutrank_perc = z-standardized percentage of group members outranked, scaleage = z-standardized age; post_hurricane1 = after the hurricane; sex:M = males; season:rainy = wet season. Model included a random effect for animal id.

**Table S8.** Intensity of *B. coli* infection as a function of social status (main effect).

| | Infection risk | | |
|---|---|---|---|
| *Predictors* | *Log-Mean* | *std. Error* | *CI (89%)* |
| Intercept | 1.95 | 0.62 | 0.95 – 2.94 |
| scaleoutrank_perc | 0.15 | 0.19 | -0.17 – 0.46 |
| post_hurricane: post_hurricane1 | -0.84 | 0.49 | -1.62 – -0.04 |
| scaleage | 0.28 | 0.18 | -0.01 – 0.57 |
| sex: M | 0.20 | 0.39 | -0.42 – 0.83 |
| season: rainy | 0.08 | 0.58 | -0.86 – 1.00 |
| $N_{id}$ | 60 | | |
| Observations | 107 | | |

scaleoutrank_perc = z-standardized percentage of group members outranked, scaleage = z-standardized age; post_hurricane1 = after the hurricane; sex:M = males; season:rainy = wet season. Model included a random effect for animal id. Analysis only included animals with count above zero (zero truncated) and was right censored (count = 60).

**Table S9.** Prevalence of *S. fuelleborni* as a function of social status (main effect).

| | Infection risk | | |
|---|---|---|---|
| *Predictors* | *Log-Odds* | *std. Error* | *CI (89%)* |
| Intercept | -5.42 | 1.84 | -8.90 – -2.85 |
| scaleoutrank_perc | 0.33 | 0.60 | -0.65 – 1.38 |
| scaleage | -0.26 | 0.60 | -1.31 – 0.73 |
| sex: M | 1.11 | 1.12 | -0.69 – 3.10 |
| post_hurricane: post_hurricane1 | 1.94 | 1.24 | 0.03 – 4.14 |
| season: rainy | -0.14 | 1.33 | -2.36 – 2.04 |
| $N_{id}$ | 100 | | |
| Observations | 199 | | |

scaleoutrank_perc = z-standardized percentage of group members outranked, scaleage = z-standardized age; post_hurricane1 = after the hurricane; sex:M = males; season:rainy = wet season. Model included a random effect for animal id.

**Table S10.** Prevalence of *T. trichiura* infection as a function of social status (main effect).

| | Infection risk | | |
|---|---|---|---|
| *Predictors* | *Log-Odds* | *std. Error* | *CI (89%)* |
| Intercept | -4.68 | 1.58 | -7.63 – -2.46 |
| scaleoutrank_perc | -0.22 | 0.52 | -1.12 – 0.62 |
| scaleage | 0.43 | 0.50 | -0.35 – 1.31 |
| sex: M | 1.11 | 0.98 | -0.41 – 2.81 |
| post_hurricane: post_hurricane1 | 2.09 | 1.09 | 0.46 – 4.04 |
| season: rainy | -0.47 | 1.15 | -2.44 – 1.34 |
| N id | 100 | | |
| Observations | 199 | | |

scaleoutrank_perc = z-standardized percentage of group members outranked, scaleage = z-standardized age; post_hurricane1 = after the hurricane; sex:M = males; season:rainy = wet season. Model included a random effect for animal id.

**Table S11.** Intensity of nematode infection as a function of social status (main effect).

| | Infection risk | | |
|---|---|---|---|
| *Predictors* | *Log-Mean* | *std. Error* | *CI (89%)* |
| Intercept | 0.67 | 0.61 | -0.31 – 1.66 |
| scaleoutrank_perc | 0.03 | 0.23 | -0.35 – 0.40 |
| post_hurricane: post_hurricane1 | 0.46 | 0.46 | -0.30 – 1.21 |
| sex: M | -0.26 | 0.42 | -0.93 – 0.43 |
| scaleage | -0.19 | 0.21 | -0.53 – 0.16 |
| season: rainy | 0.43 | 0.49 | -0.38 – 1.20 |
| N id | 35 | | |
| Observations | 52 | | |

scaleoutrank_perc = z-standardized percentage of group members outranked, scaleage = z-standardized age; post_hurricane1 = after the hurricane; sex:M = males; season:rainy = wet season. Model included a random effect for animal id. Analysis only included animals with count above zero (zero truncated).

**Table S11.** Prevalence of *B. coli* infection pre and post hurricane as a function of social status.

| | Infection risk | | |
|---|---|---|---|
| *Predictors* | *Log-Odds* | *std. Error* | *CI (89%)* |
| Intercept | 3.04 | 1.93 | -0.06 – 6.38 |
| scaleoutrank_perc | 0.40 | 0.87 | -1.06 – 1.85 |
| post_hurricane | -3.51 | 1.61 | -6.42 – -1.07 |
| scaleage | 0.52 | 0.75 | -0.67 – 1.85 |
| sex: M | 0.52 | 1.38 | -1.77 – 2.83 |
| season: rainy | -1.42 | 1.70 | -4.28 – 1.25 |
| scaleoutrank_perc:post_hurricane | -1.55 | 1.32 | -3.83 – 0.55 |
| $N_{id}$ | 100 | | |
| Observations | 199 | | |

scaleoutrank_perc = z-standardized percentage of group members outranked, scaleage = z-standardized age; post_hurricane1 = after the hurricane; sex:M = males; season:rainy = wet season. Model included a random effect for animal id.

**Table S12.** Intensity of *B. coli* infection pre and post hurricane as a function of social status.

| | Infection risk | | |
|---|---|---|---|
| *Predictors* | *Log-Mean* | *std. Error* | *CI (89%)* |
| Intercept | 1.95 | 0.63 | 0.94 – 2.96 |
| scaleoutrank_perc | 0.07 | 0.24 | -0.31 – 0.45 |
| post_hurricane | -0.79 | 0.50 | -1.59 – 0.02 |
| scaleage | 0.27 | 0.18 | -0.02 – 0.57 |
| sex: M | 0.16 | 0.40 | -0.48 – 0.80 |
| season: rainy | 0.09 | 0.59 | -0.86 – 1.05 |
| scaleoutrank_perc:post_hurricane | 0.24 | 0.39 | -0.39 – 0.86 |
| $N_{id}$ | 60 | | |
| Observations | 107 | | |

scaleoutrank_perc = z-standardized percentage of group members outranked, scaleage = z-standardized age; post_hurricane1 = after the hurricane; sex:M = males; season:rainy = wet season. Model included a random effect for animal id. Analysis only included animals with count above zero (zero truncated) and was right censored (count = 60).

**Table S13.** Prevalence of *S. fuelleborni* infection pre and post hurricane as a function of social status.

| | Infection risk | | |
|---|---|---|---|
| *Predictors* | *Log-Odds* | *std. Error* | *CI (89%)* |
| Intercept | -5.56 | 1.93 | -9.14 – -2.89 |
| scaleoutrank_perc | 0.14 | 0.80 | -1.17 – 1.50 |
| post_hurricane | 1.95 | 1.25 | 0.04 – 4.20 |
| scaleage | -0.27 | 0.61 | -1.36 – 0.75 |
| sex: M | 1.03 | 1.18 | -0.87 – 3.09 |
| season: rainy | -0.12 | 1.35 | -2.37 – 2.10 |
| scaleoutrank_perc:post_hurricane | 0.48 | 1.12 | -1.32 – 2.36 |
| $N_{id}$ | 100 | | |
| Observations | 199 | | |

scaleoutrank_perc = z-standardized percentage of group members outranked, scaleage = z-standardized age; post_hurricane1 = after the hurricane; sex:M = males; season:rainy = wet season. Model included a random effect for animal id.

**Table S14.** Prevalence of *T. trichiura* infection pre and post hurricane as a function of social status.

| | Infection risk | | |
|---|---|---|---|
| *Predictors* | *Log-Odds* | *std. Error* | *CI (89%)* |
| Intercept | -5.17 | 1.19 | -7.58 – -3.53 |
| post_hurricane | 2.27 | 0.99 | 0.81 – 4.11 |
| scaleoutrank_perc | -0.06 | 0.68 | -1.21 – 1.09 |
| scaleage | 0.46 | 0.48 | -0.32 – 1.31 |
| sex: M | 1.16 | 1.01 | -0.43 – 2.88 |
| post_hurricane:scaleoutrank_perc | -0.30 | 0.95 | -1.90 – 1.26 |
| $N_{id}$ | 100 | | |
| Observations | 199 | | |

scaleoutrank_perc = z-standardized percentage of group members outranked, scaleage = z-standardized age; post_hurricane1 = after the hurricane; sex:M = males; season:rainy = wet season. Model included a random effect for animal id.

**Table S15.** Intensity of nematode infection pre and post hurricane as a function of social status.

| | Infection risk | | |
|---|---|---|---|
| *Predictors* | *Log-Mean* | *std. Error* | *CI (89%)* |
| Intercept | 0.66 | 0.61 | -0.34 – 1.68 |
| scaleoutrank_perc | 0.04 | 0.44 | -0.66 – 0.76 |
| post_hurricane | 0.47 | 0.47 | -0.30 – 1.24 |
| sex: M | -0.25 | 0.44 | -0.99 – 0.47 |
| scaleage | -0.19 | 0.22 | -0.56 – 0.17 |
| season: rainy | 0.43 | 0.50 | -0.41 – 1.23 |
| scaleoutrank_perc:post_hurricane | -0.02 | 0.52 | -0.87 – 0.81 |
| N $_{id}$ | 35 | | |
| Observations | 52 | | |

scaleoutrank_perc = z-standardized percentage of group members outranked, scaleage = z-standardized age; post_hurricane1 = after the hurricane; sex:M = males; season:rainy = wet season. Model included a random effect for animal id. Analysis only included animals with count above zero (zero truncated).

**Table S16.** Prevalence of *B. coli* infection as a function of the number of weak connections in the grooming network.

| | Infection risk | | |
|---|---|---|---|
| *Predictors* | *Log-Odds* | *std. Error* | *CI (89%)* |
| Intercept | 0.32 | 1.11 | -1.53 – 2.22 |
| scalestd_weak_groom | -1.71 | 1.39 | -4.19 – 0.46 |
| scaleperc_rank | 0.64 | 0.96 | -0.92 – 2.36 |
| scaleage | 1.77 | 1.08 | 0.16 – 3.83 |
| sex: M | -0.10 | 1.64 | -2.87 – 2.63 |
| N $_{id}$ | 70 | | |
| Observations | 140 | | |

scalestd_weak_groom = z-standardised number of weak grooming connections (previously standardised within groups), scaleoutrank_perc = z-standardized percentage of group members outranked, scaleage = z-standardized age; sex:M = males. Model included a random effect for animal id.

**Table S17.** Prevalence of *B. coli* infection as a function of the number of weak connections in the proximity network.

| | Infection risk | | |
|---|---|---|---|
| *Predictors* | *Log-Odds* | *std. Error* | *CI (89%)* |
| Intercept | 0.56 | 1.07 | -1.24 – 2.41 |
| scalestd_weak_prox | -2.03 | 1.18 | -4.16 – -0.18 |
| scaleperc_rank | 0.94 | 0.93 | -0.56 – 2.63 |
| scaleage | 2.16 | 1.10 | 0.55 – 4.26 |
| sex: M | -0.66 | 1.64 | -3.46 – 2.05 |
| N id | 70 | | |
| Observations | 140 | | |

scalestd_weak_prox = z-standardised number of weak proximity connections (previously standardised within groups), scaleoutrank_perc = z-standardized percentage of group members outranked, scaleage = z-standardized age; sex:M = males. Model included a random effect for animal id.

**Table S18.** Prevalence of *S. fuelleborni* infection as a function of the number of weak connections in the grooming network.

| | Infection risk | | |
|---|---|---|---|
| *Predictors* | *Log-Odds* | *std. Error* | *CI (89%)* |
| Intercept | -4.46 | 1.40 | -7.50 – -2.68 |
| scalestd_weak_groom | 0.32 | 0.88 | -1.14 – 1.85 |
| scaleperc_rank | -0.13 | 0.72 | -1.39 – 1.11 |
| scaleage | -0.28 | 0.75 | -1.61 – 0.96 |
| sex: M | -0.57 | 1.33 | -2.89 – 1.62 |
| N id | 70 | | |
| Observations | 140 | | |

scalestd_weak_groom = z-standardised number of weak grooming connections (previously standardised within groups), scaleoutrank_perc = z-standardized percentage of group members outranked, scaleage = z-standardized age; sex:M = males. Model included a random effect for animal id.

**Table S19.** Prevalence of *S. fuelleborni* infection as a function of the number of weak connections in the proximity network.

| | Infection risk | | |
|---|---|---|---|
| *Predictors* | *Log-Odds* | *std. Error* | *CI (89%)* |
| Intercept | -4.53 | 1.40 | -7.51 – -2.75 |
| scalestd_weak_prox | 0.71 | 0.86 | -0.66 – 2.26 |
| scaleperc_rank | -0.26 | 0.74 | -1.55 – 0.98 |
| scaleage | -0.45 | 0.78 | -1.85 – 0.82 |
| sex: M | -0.34 | 1.36 | -2.66 – 1.89 |
| N id | 70 | | |
| Observations | 140 | | |

scalestd_weak_prox = z-standardised number of weak proximity connections (previously standardised within groups), scaleoutrank_perc = z-standardized percentage of group members outranked, scaleage = z-standardized age; sex:M = males. Model included a random effect for animal id.

**Table S20.** Prevalence of *T. trichiura* infection as a function of the number of weak connections in the grooming network.

| | Infection risk | | |
|---|---|---|---|
| *Predictors* | *Log-Odds* | *std. Error* | *CI (89%)* |
| Intercept | -4.47 | 1.40 | -7.47 – -2.69 |
| scalestd_weak_groom | 0.32 | 0.88 | -1.14 – 1.83 |
| scaleperc_rank | -0.12 | 0.73 | -1.40 – 1.11 |
| scaleage | -0.28 | 0.75 | -1.62 – 0.95 |
| sex: M | -0.56 | 1.33 | -2.88 – 1.61 |
| N id | 70 | | |
| Observations | 140 | | |

scalestd_weak_groom = z-standardised number of weak grooming connections (previously standardised within groups), scaleoutrank_perc = z-standardized percentage of group members outranked, scaleage = z-standardized age; sex:M = males. Model included a random effect for animal id.

**Table S21.** Prevalence of *T. trichiura* infection as a function of the number of weak connections in the proximity network.

| | Infection risk | | |
|---|---|---|---|
| *Predictors* | *Log-Odds* | *std. Error* | *CI (89%)* |
| Intercept | -4.54 | 1.41 | -7.58 – -2.76 |
| scalestd_weak_prox | 0.71 | 0.86 | -0.66 – 2.27 |
| scaleperc_rank | -0.27 | 0.74 | -1.57 – 0.98 |
| scaleage | -0.45 | 0.79 | -1.86 – 0.83 |
| sex: M | -0.33 | 1.36 | -2.67 – 1.91 |
| N id | 70 | | |
| Observations | 140 | | |

scalestd_weak_prox = z-standardised number of weak proximity connections (previously standardised within groups), scaleoutrank_perc = z-standardized percentage of group members outranked, scaleage = z-standardized age; sex:M = males. Model included a random effect for animal id.

**Table S22.** Univariate model to test effect of sex on *S. fuelleborni* infection risk.

| | Infection risk | | |
|---|---|---|---|
| *Predictors* | *Log-Odds* | *std. Error* | *CI (89%)* |
| Intercept | -4.03 | 0.99 | -6.00 – -2.72 |
| sex: M | 1.12 | 0.93 | -0.39 – 2.72 |
| N id | 100 | | |
| Observations | 199 | | |

sex:M = males. Model included a random effect for animal id.

**Table S23.** Univariate model to test effect of age on *S. fuelleborni* infection risk.

| | Infection risk | | |
|---|---|---|---|
| *Predictors* | *Log-Odds* | *std. Error* | *CI (89%)* |
| Intercept | -3.72 | 0.94 | -5.62 – -2.55 |
| scaleage | -0.06 | 0.53 | -0.94 – 0.83 |
| N id | 100 | | |
| Observations | 199 | | |

scaleage = z-standardized age. Model included a random effect for animal id.

**Table S24.** Univariate model to test effect of rank on *S. fuelleborni* infection risk.

| | Infection risk | | |
|---|---|---|---|
| *Predictors* | *Log-Odds* | *std. Error* | *CI (89%)* |
| Intercept | -3.62 | 0.89 | -5.52 – -2.55 |
| scaleoutrank_perc | 0.32 | 0.51 | -0.51 – 1.16 |
| N id | 100 | | |
| Observations | 199 | | |

scaleoutrank_perc = z-standardized percentage of group members outranked. Model included a random effect for animal id.

**Table S25.** Univariate model to test effect of sex on *T. trichiura* infection risk.

| | Infection risk | | |
|---|---|---|---|
| *Predictors* | *Log-Odds* | *std. Error* | *CI (89%)* |
| Intercept | -3.62 | 0.86 | -5.33 – -2.47 |
| sex: M | 0.73 | 0.90 | -0.67 – 2.31 |
| N id | 100 | | |
| Observations | 199 | | |

sex:M = males. Model included a random effect for animal id.

**Table S26.** Univariate model to test effect of age on *T. trichiura* infection risk.

| | Infection risk | | |
|---|---|---|---|
| *Predictors* | *Log-Odds* | *std. Error* | *CI (89%)* |
| Intercept | -3.27 | 0.72 | -4.90 – -2.29 |
| scaleage | 0.54 | 0.43 | -0.11 – 1.33 |
| N id | 100 | | |
| Observations | 199 | | |

scaleage = z-standardized age. Model included a random effect for animal id.

**Table S27.** Univariate model to test effect of rank on *T. trichiura* infection risk.

| | Infection risk | | |
|---|---|---|---|
| *Predictors* | *Log-Odds* | *std. Error* | *CI (89%)* |
| Intercept | -3.39 | 0.80 | -5.12 – -2.39 |
| scaleoutrank_perc | -0.11 | 0.44 | -0.94 – 0.65 |
| N id | 100 | | |
| Observations | 199 | | |

scaleoutrank_perc = z-standardized percentage of group members outranked. Model included a random effect for animal id.

**Table S28.** Prevalence of *S. fuelleborni* infection as a function of the number of weak partners in the grooming network (univariate model).

| | Infection risk | | |
|---|---|---|---|
| *Predictors* | *Log-Odds* | *std. Error* | *CI (89%)* |
| Intercept | -3.93 | 1.08 | -6.27 – -2.55 |
| scalestd_weak_groom | 0.26 | 0.73 | -0.94 – 1.52 |
| N id | 70 | | |
| Observations | 140 | | |

scalestd_weak_groom = z-standardised number of weak grooming connections (previously standardised within groups), Model included a random effect for animal id.

**Table S29.** Prevalence of *S. fuelleborni* infection as a function of the number of weak partners in the proximity network (univariate model).

| | Infection risk | | |
|---|---|---|---|
| *Predictors* | *Log-Odds* | *std. Error* | *CI (89%)* |
| Intercept | -3.94 | 1.08 | -6.27 – -2.56 |
| scalestd_weak_prox | 0.48 | 0.66 | -0.56 – 1.67 |
| N id | 70 | | |
| Observations | 140 | | |

scalestd_weak_prox = z-standardised number of weak proximity connections (previously standardised within groups), Model included a random effect for animal id.

**Table S30.** Prevalence of *T. trichiura* infection as a function of the number of weak partners in the grooming network (univariate model).

| | **Infection risk** | | |
|---|---|---|---|
| *Predictors* | *Log-Odds* | *std. Error* | *CI (89%)* |
| Intercept | -3.60 | 0.95 | -5.63 – -2.37 |
| scalestd_weak_groom | 0.04 | 0.64 | -1.08 – 1.11 |
| N id | 70 | | |
| Observations | 140 | | |

scalestd_weak_groom = z-standardised number of weak grooming connections (previously standardised within groups), Model included a random effect for animal id.

**Table S31.** Prevalence of *T. trichiura* infection as a function of the number of weak partners in the proximity network (univariate model).

| | **Infection risk** | | |
|---|---|---|---|
| *Predictors* | *Log-Odds* | *std. Error* | *CI (89%)* |
| Intercept | -3.57 | 0.95 | -5.58 – -2.36 |
| scalestd_weak_prox | 0.37 | 0.56 | -0.56 – 1.35 |
| N id | 70 | | |
| Observations | 140 | | |

scalestd_weak_prox = z-standardised number of weak proximity connections (previously standardised within groups), Model included a random effect for animal id.

**Table S32.** Prevalence of *B. coli* infection as a function of the strength to strong connections in the grooming network.

| | **Infection risk** | | |
|---|---|---|---|
| *Predictors* | *Log-Odds* | *std. Error* | *CI (89%)* |
| Intercept | 0.28 | 1.13 | -1.61 – 2.23 |
| scalestd_topstr_groom | -1.06 | 1.41 | -3.47 – 1.24 |
| scaleperc_rank | 0.66 | 0.99 | -0.95 – 2.44 |
| scaleage | 1.77 | 1.10 | 0.14 – 3.88 |
| sex: M | 0.04 | 1.66 | -2.77 – 2.83 |
| $N_{id}$ | 70 | | |
| Observations | 140 | | |

scalestd_topstr_groom = z-standardised strength to strong grooming connections (previously standardised within groups), scaleoutrank_perc = z-standardized percentage of group members outranked, scaleage = z-standardized age; sex:M = males. Model included a random effect for animal id.

**Table S33.** Prevalence of *B. coli* infection as a function of the strength to strong connections in the proximity network.

| | **Infection risk** | | |
|---|---|---|---|
| *Predictors* | *Log-Odds* | *std. Error* | *CI (89%)* |
| Intercept | 0.71 | 1.08 | -1.10 – 2.53 |
| scalestd_topstr_prox | -2.07 | 1.29 | -4.36 – -0.02 |
| scaleperc_rank | 1.39 | 1.04 | -0.28 – 3.21 |
| scaleage | 2.12 | 1.08 | 0.54 – 4.19 |
| sex: M | -1.04 | 1.73 | -3.97 – 1.81 |
| $N_{id}$ | 70 | | |
| Observations | 140 | | |

scalestd_topstr_prox = z-standardised strength to strong proximity connections (previously standardised within groups), scaleoutrank_perc = z-standardized percentage of group members outranked, scaleage = z-standardized age; sex:M = males. Model included a random effect for animal id.

**Table S34.** Prevalence of *S. fuelleborni* infection as a function of the strength to strong connections in the grooming network.

| | **Infection risk** | | |
|---|---|---|---|
| *Predictors* | *Log-Odds* | *std. Error* | *CI (89%)* |
| Intercept | -4.42 | 1.39 | -7.44 – -2.67 |
| scalestd_topstr_groom | 0.05 | 0.96 | -1.65 – 1.63 |
| scaleperc_rank | -0.12 | 0.73 | -1.38 – 1.11 |
| scaleage | -0.27 | 0.74 | -1.58 – 0.97 |
| sex: M | -0.61 | 1.33 | -2.90 – 1.55 |
| $N_{id}$ | 70 | | |
| Observations | 140 | | |

scalestd_topstr_groom = z-standardised strength to strong grooming connections (previously standardised within groups), scaleoutrank_perc = z-standardized percentage of group members outranked, scaleage = z-standardized age; sex:M = males. Model included a random effect for animal id.

**Table S35.** Prevalence of *S. fuelleborni* infection as a function of the strength to strong connections in the proximity network.

| | **Infection risk** | | |
|---|---|---|---|
| *Predictors* | *Log-Odds* | *std. Error* | *CI (89%)* |
| Intercept | -4.54 | 1.41 | -7.53 – -2.73 |
| scalestd_topstr_prox | 0.49 | 0.99 | -1.11 – 2.20 |
| scaleperc_rank | -0.34 | 0.84 | -1.81 – 1.04 |
| scaleage | -0.39 | 0.78 | -1.80 – 0.88 |
| sex: M | -0.31 | 1.44 | -2.78 – 2.06 |
| $N_{id}$ | 70 | | |
| Observations | 140 | | |

scalestd_topstr_prox = z-standardised strength to strong proximity connections (previously standardised within groups), scaleoutrank_perc = z-standardized percentage of group members outranked, scaleage = z-standardized age; sex:M = males. Model included a random effect for animal id.

**Table S36.** Prevalence of *T. trichiura* infection as a function of the strength to strong connections in the grooming network.

| | **Infection risk** | | |
|---|---|---|---|
| *Predictors* | *Log-Odds* | *std. Error* | *CI (89%)* |
| Intercept | -4.43 | 1.39 | -7.44 – -2.68 |
| scalestd_topstr_groom | 0.05 | 0.96 | -1.64 – 1.63 |
| scaleperc_rank | -0.12 | 0.73 | -1.39 – 1.11 |
| scaleage | -0.27 | 0.74 | -1.59 – 0.96 |
| sex: M | -0.61 | 1.34 | -2.91 – 1.58 |
| $N_{id}$ | 70 | | |
| Observations | 140 | | |

scalestd_topstr_groom = z-standardised strength to strong grooming connections (previously standardised within groups), scaleoutrank_perc = z-standardized percentage of group members outranked, scaleage = z-standardized age; sex:M = males. Model included a random effect for animal id.

**Table S37.** Prevalence of *T. trichiura* infection as a function of the strength to strong connections in the proximity network.

| | **Infection risk** | | |
|---|---|---|---|
| *Predictors* | *Log-Odds* | *std. Error* | *CI (89%)* |
| Intercept | -4.34 | 1.27 | -7.05 – -2.69 |
| scalestd_topstr_prox | 0.40 | 0.79 | -0.90 – 1.71 |
| scaleperc_rank | -0.27 | 0.71 | -1.46 – 0.91 |
| scaleage | -0.31 | 0.69 | -1.48 – 0.81 |
| sex: M | -0.24 | 1.10 | -2.03 – 1.54 |
| $N_{id}$ | 70 | | |
| Observations | 140 | | |

scalestd_topstr_prox = z-standardised strength to strong proximity connections (previously standardised within groups), scaleoutrank_perc = z-standardized percentage of group members outranked, scaleage = z-standardized age; sex:M = males. Model included a random effect for animal id.

**Table S38.** Prevalence of *S. fuelleborni* infection as a function of the strength to strong connections in the grooming network (univariate model).

| | Infection risk | | |
|---|---|---|---|
| *Predictors* | *Log-Odds* | *std. Error* | *CI (89%)* |
| Intercept | -3.90 | 1.07 | -6.20 – -2.54 |
| scalestd_topstr_groom | 0.01 | 0.79 | -1.40 – 1.32 |
| N id | 70 | | |
| Observations | 140 | | |

scalestd_topstr_groom = z-standardised strength to strong grooming connections (previously standardised within groups), Model included a random effect for animal id.

**Table S39.** Prevalence of *S. fuelleborni* infection as a function of the strength to strong connections in the proximity network (univariate model).

| | Infection risk | | |
|---|---|---|---|
| *Predictors* | *Log-Odds* | *std. Error* | *CI (89%)* |
| Intercept | -3.93 | 1.08 | -6.26 – -2.56 |
| scalestd_topstr_prox | 0.23 | 0.64 | -0.84 – 1.36 |
| N id | 70 | | |
| Observations | 140 | | |

scalestd_topstr_prox = z-standardised strength to strong proximity connections (previously standardised within groups), Model included a random effect for animal id.

**Table S40.** Prevalence of *T. trichiura* infection as a function of the strength to strong connections in the grooming network (univariate model).

| | Infection risk | | |
|---|---|---|---|
| *Predictors* | *Log-Odds* | *std. Error* | *CI (89%)* |
| Intercept | -3.58 | 0.95 | -5.59 – -2.37 |
| scalestd_topstr_groom | 0.20 | 0.64 | -0.87 – 1.29 |
| N id | 70 | | |
| Observations | 140 | | |

scalestd_topstr_groom = z-standardised strength to strong grooming connections (previously standardised within groups), Model included a random effect for animal id.

**Table S41.** Prevalence of *T. trichiura* infection as a function of the strength to strong connections in the proximity network (univariate model).

| | Infection risk | | |
|---|---|---|---|
| *Predictors* | *Log-Odds* | *std. Error* | *CI (89%)* |
| Intercept | -3.58 | 0.95 | -5.61 – -2.35 |
| scalestd_topstr_prox | 0.25 | 0.56 | -0.72 – 1.19 |
| $N_{id}$ | 70 | | |
| Observations | 140 | | |

scalestd_topstr_prox= z-standardised strength to strong proximity connections (previously standardised within groups), Model included a random effect for animal id.
